# Agent-based modeling of tumor-immune interactions reveals determinants of final tumor states

**DOI:** 10.1101/2023.09.06.556617

**Authors:** Manal Ahmidouch, Neel Tangella, Stacey D. Finley

## Abstract

Interactions between tumor and immune cells in the tumor microenvironment (TME) influence tumor growth and the tumor’s response to treatment. Excitingly, this complex landscape of tumor-immune interactions can be studied using computational modeling. Mathematical oncology can provide quantitative insights into the TME, serving as a framework for understanding tumor dynamics. Here, we use an agent-based model to simulate the interactions among cancer cells, macrophages (naïve, M1, and M2), and T cells (active CD8+ and inactive) in a 2D representation of the TME. Key diffusible factors, IL-4 and IFN-γ, are also incorporated. We apply the model to predict how cell-specific properties influence tumor progression. The model predictions and analyses revealed the relationships between different cell populations and highlighted the importance of macrophages and T cells in shaping the TME. Thus, we quantify how components of the TME influence the final tumor state and the effects of macrophage-based therapies. The findings emphasize the significant role of computational models in unraveling the intricate dynamics of tumor-immune interactions and their potential for guiding the development of tailored immunotherapeutic strategies. This study provides a foundation for future investigations aiming to refine and expand the model, validate predictions experimentally, and pave the way for improved cancer treatments.

## Introduction

As cancer immunotherapies demonstrate success in certain tumor settings and amass attention^1^, creating models to simulate the effects of these treatment becomes increasingly crucial. In particular, mathematical models of tumor growth are able to account for the complex and adaptive interactions between tumor and immune cells^2^. Immune cells have the innate ability to detect malignant tumor cells as “not self” and destroy them. Tumor cells, however, are able to suppress this immune function and alter immune cells such that they assist in tumor growth, rather than diminish it^3^.The altered immune cells also suppress the immune system^4^. Thus, significant effort has been focused on developing immunotherapies that overcome such tumor-promoting interactions^5^.

Macrophage-targeted therapies hold great promise due to the vital role macrophages play in tumor progression^6^. There are two broad classes of macrophages identified using *in vitro* studies, termed M1 and M2^7^. M1 macrophages are able to activate neighboring T cells. In contrast, M2 macrophages prevent neighboring CD8+ T cells from becoming activated by secreting inhibitory cytokines such as interleukin-4 (IL-4). Tumor cells also secrete IL-4, contributing to the anti-tumor effect of macrophages^8^. When the T cells become fully activated, they proliferate and kill the nearby cancer cells. They also secrete interferon-γ (IFN-γ), which attracts M1 macrophages to the tumor site and further promotes M1 activity^9^. In sum, M1 macrophages promote T cell activation to produce CD8+ T cells and M2 macrophages inhibit it, with the CD8+ T cells destroying the tumor cells via cytolytic activity^10^.

Many current immunotherapies target immunosuppressive mechanisms. However, since tumor-immune interactions are complex, the result of these immunotherapies can differ from what is expected when studied using experimental models^11^. Additionally, given the intricacy of the tumor microenvironment (TME), *in vitro* and *in vivo* studies alone cannot resolve the potential effects of immunotherapy. Excitingly, the tumor and immune cell interactions can be better understood using computational models^2,12^. Computational models allow us to simulate a plethora of immune cell interactions, in less time and with fewer resources than with a purely experimental approach. Importantly, the information used in these computational models is based on previous biological experiments, so the models can be used to make reliable and physiologically-relevant predictions of tumor behavior^13^. While many computational models simulate individual interactions within the TME, it is ideal to create a detailed model to more fully represent the TME and the immune cell interactions involved. In particular, agent-based models (ABMs) combine individual interactions into a complex system of cellular behaviors, making them ideal for modeling the TME^14^. ABMs are commonly utilized to capture the spatial characteristics of a system. These models consist of individual agents representing cells that interact with their neighbors based on experiment-derived rules that reflect the cells’ biological roles^15^. By following these rules, the agents behave in a way that captures the overall behavior of the system.

In this study, we employ an ABM to simulate the early growth of tumors in a 2D representation, resembling a tumor slice^16^. Our model incorporates three primary cell types: T cells, macrophages, and cancer cells. Additionally, we consider the production and diffusion of diffusible factors using partial differential equations. The parameter values used in the model are derived from previous modeling efforts or are based on experimental observations. By varying these parameters, we can simulate the dynamic tumor behavior, ultimately establishing whether a tumor progresses towards a pro-tumor state or an anti-tumor end state. By investigating the intricate dynamics of tumor-immune cell interactions through our ABM, we aim to shed light on the underlying mechanisms governing tumor behavior and response to immunotherapy. In particular, we uncover relationships between subsets of cells and how these relationships evolve over time to shape tumor growth. We quantify how specific cell types at discrete times influence the final tumor state. Overall, our work provides a more detailed understanding of how distinct cell types and their interactions contribute to tumor progression.

## Methods

### Model Overview

An ABM was employed to simulate the interactions among diverse cell populations within the TME. The model represents the early stages of tumor development and incorporates three primary cell types: cancer cells, macrophages, and T cells. The model also incorporates two diffusible factors, IL-4 and IFN-γ, which play crucial roles in influencing the immune state of the TME. The simulation takes place in a two-dimensional environment, comprised of a 100-by-100 grid representing a 1.5 mm^2^ tissue slice. Thus, each site within the grid is 1.5 μm^2^. As this is approximately the diameter of cells, each site can be occupied by only one cell at a time. Detailed information about the ABM used in this study can be found in^16^ and accessed at the GitHub repository here: https://github.com/FinleyLabUSC/Early-TME-ABM-PLOS-Comp-Bio.

### Cell Populations

The primary cell types included in the model are cancer cells, macrophages (naïve, M1, and M2), and T cells (active CD8+ and inactive). M1 macrophages are able to activate neighboring T cells, while M2 macrophages inhibit T cells from becoming active. Cancer cells are the main contributors to tumor growth, while macrophages and T cells interact with the cancer cells to influence tumor growth. It is important to note that this model aims to understand generalized tumor behavior rather than focusing on a specific tumor type. The simulations were first performed without the application of any treatment.

Macrophages initially enter the microenvironment as naïve M0 macrophages, and their polarization into the M1 or M2 state is determined by the presence of diffusible factors in the local environment at a given time. IL-4, secreted by the tumor cells and M2 macrophages, promotes a pro-tumor state. In comparison, IFN-γ, secreted by activated T cells, induces M1 differentiation, thereby fostering an anti-tumor environment. Macrophages sense the concentrations of IL-4 and IFN-γ in their local microenvironment and process these input signals via an intracellular signaling model encoded as a trained neural network, resulting in differentiation. Macrophages can re-differentiate every 24 hours based on the local diffusible factors.

The recruitment of T cells into the TME is crucial for tumor eradication. In the model, T cells possess the ability to migrate towards the tumor and undergo full activation upon encountering a cancer cell. Once fully activated, T cells can proliferate and effectively eliminate neighboring cancer cells. Additionally, fully active T cells secrete IFN-γ, which influences the differentiation of naïve macrophages into the M1 macrophage state, enhancing the anti-tumor environment.

### Simulation of Diffusible Factors

To simulate the production and diffusion of IL-4 and IFN-γ, the model incorporates partial differential equations that describe the spatiotemporal evolution of the diffusible factors. Parameters for these equations were derived from previous modeling efforts or experimental observations.

### Parameter Alteration and Simulation Design

To investigate the impact of specific parameters on the TME, we conducted simulations over a 200-day period. Each simulation was repeated 100 times to account for the inherent stochasticity of ABMs, and the resulting data were averaged at each time point for analysis. The key parameters under investigation were the cancer cell cycle length, macrophage recruitment rate, and T cell recruitment rate. We explored cell cycle lengths of 20, 25, and 30 hours, T cell recruitment rates of 1, 5, and 10 cells/hour and macrophage recruitment rates of 1×10^-9^ and 1×10^-8^ cells/site/second.

### Outcome Measures

The primary outcome measure assessed in this study was the number of cancer cells at each time point throughout the simulation. This served as a quantifiable indicator of tumor growth. By monitoring the changes in cancer cell population size over time, we gained insights into the dynamics of tumor growth. Additionally, the dynamic population sizes of different cell types within the TME were analyzed to gain a comprehensive understanding of their relationships.

The behaviors of macrophages (naïve, M1, and M2) and T cells (inactive and active CD8) were studied to evaluate their influence on the TME and potential roles in tumor progression or suppression.

In addition to studying temporal dynamics, we also examined the end state for each cell type. The end state refers to the final population size of a given cell type at the conclusion of the simulation. By identifying and analyzing the end states, we gained valuable insights into the regulatory mechanisms within the TME. This analysis allowed us to characterize the final configurations of cell populations and understand the interplay between different cell types.

### Data Analysis

Data visualization techniques were employed to depict the population size trends of cell types and to compare the impact of different parameter settings on tumor growth.

### Partial Least Squares Analysis

A common technique used to perform regression on high-dimensional data is partial least-squares (PLS). Partial least squares regression (PLSR) aims to reduce the dimensionality of the data by constructing latent variables that represent the inputs. In our study, we employ a modified version of PLS called partial least squares discriminant analysis (PLS-DA). PLS-DA is a supervised learning approach that is particularly suitable for modeling relationships between species’ concentrations and their contributions to different simulation end states. PLS-DA as a method for understanding feature importance in high-dimensional datasets has seen increasing use in the field of metabolomics^17^. The inputs to the PLS-DA model (also called “predictors”) are the number of cellular and molecular species of each type at discrete time points. For the output, we define two categories representing favorable and unfavorable TME end states, allowing us to perform PLS-DA and examine the features of the TME that influence the model’s outcome. Here, a favorable end state is when the number of active CD8+ T cells exceeds the number of cancer cells. In contrast, an unfavorable end state is when the number of cancer cells exceeds the number of active CD8+ T cells.

### Model Selection

We vary the number of components used in PLS-DA to determine the model with the highest predictive power. To select the PLS-DA model with the highest predictive power and control against PLS-DA’s tendency to overfit, we perform *k*-fold cross-validation. This technique involves splitting the input data into *k* groups, where one group serves as the test set and the remaining *k*-1 groups are used as the training set. We shuffle the data before each iteration of cross-validation to ensure equal representation. The PLS-DA model that achieves the highest accuracy on the test dataset is selected for subsequent model projection and analysis.

### VIP Score

The variable importance of projection (VIP) score is a common metric accompanying PLS-DA models. It quantifies the relative importance of each input variable to the model^18^. The VIP score is calculated as the weighted sum of squared correlations between the model components and the original predictor variable. Typically, a VIP score of 1 or greater is considered indicative of above-average importance^18^, and we use this threshold here.

### Loadings

Analyzing the loadings in a PLS-DA model is crucial for determining the direct linear contributions of input variables to the model projection. Strongly positive or negative loadings indicate that the input variable plays a more significant role in discriminating the model’s outputs compared to inputs with loadings close to zero. In our study, given the definition of the two TME end states, inputs with positive loadings drive the system towards a favorable end state, while those with negative loadings drive it towards an unfavorable end state.

By employing PLS-DA, selecting the optimal model, and analyzing VIP scores and loadings, we identify the key inputs and their contributions in driving the TME towards favorable or unfavorable end states. This analysis provides insights into the underlying mechanisms and factors influencing the TME dynamics.

## Results

### Cell-specific properties strongly influence tumor growth

The predicted dynamics observed in our simulations provide valuable insights into the relationships between different cell types within the TME. We first examined how tumor dynamics were affected by cell-specific properties, including immune cell recruitment and the length of the cancer cell cycle (**Figure 1**), as these are tumor-specific properties that will vary within the TME.

**Fig 1.**
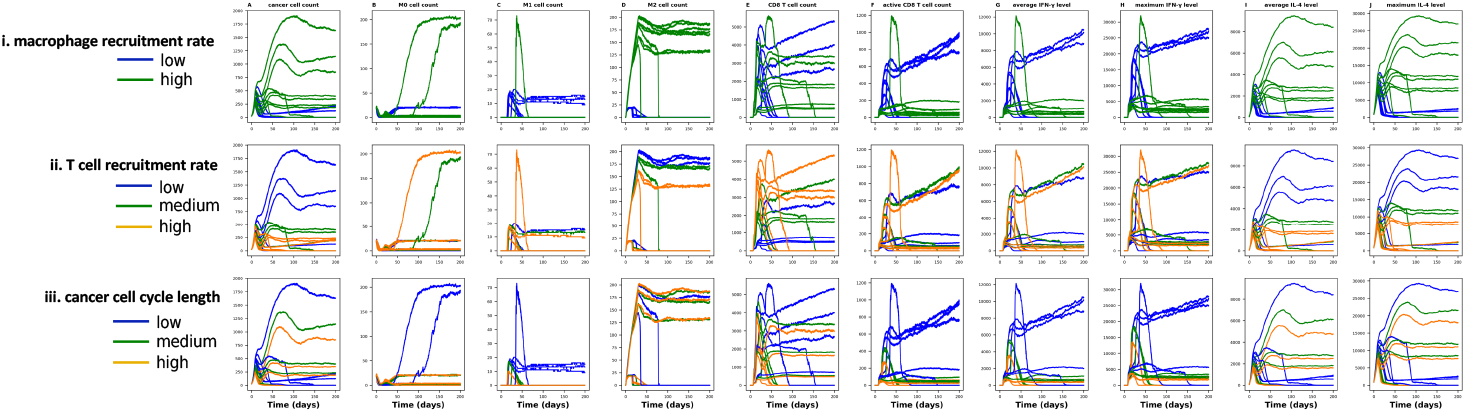
Effect of immune cell recruitment rates and cell cycle length on cell population sizes. A) cancer cell count. B) M0 cell count. C) M1 cell count. D) M2 cell count. E) CD8 T cell count. F) active CD8 T cell count. G) average IFN-γ level. H) maximum IFN-γ. I) average IL-4 level. J) maximum IL-4 level. Time courses for the 100 replicates of the model are plotted and averaged at each time point. Cell cycle lengths of 20, 25, and 30 hours, T cell recruitment rates of 1, 5, and 10 cells/hour and macrophage recruitment rates of 1×10^-8^ and 1×10^-7^ cells/site/second were explored.

Model predictions show that the balance between macrophage and T cell recruitment rates, as well as the length of the cancer cell growth cycle, influences the population sizes of different cell types and the levels of immune-related factors (**Figure 1**). Increasing the macrophage recruitment rate was found to promote a pro-tumor microenvironment, resulting in an increase in the cancer cell population (Figure 1A, row (i)). This was accompanied by an increase in the M2 macrophage population, indicating increased differentiation of naïve macrophages to the immunosuppressive state. Interestingly, the higher macrophage recruitment rate was associated with a smaller active CD8+ T cell population and lower IFN-γ levels (Figure 1F, row (i), G, row (i), and H, row (i)), suggesting a dampened anti-tumor immune response. Additionally, elevated macrophage recruitment rates led to higher IL-4 concentrations (Figure 1I, row (i) and J, row (i)), contributed by cancer cells and M2 macrophages.

Compared to increasing the macrophage recruitment rate, higher T cell recruitment contributed to an anti-tumor microenvironment, characterized by a lower number of cancer cells (Figure 1A, row (ii)). This was accompanied by fewer M2 macrophages and lower IL-4 levels (Figure 1D, row (ii), I, row (ii) and J, row (ii)). Interestingly, we did not observe a direct relationship between the size of the T cell population and the rate of T cell recruitment.

Finally, varying the length of the cancer cell growth cycle had a significant impact on the tumor microenvironment. In general, shorter cell cycle lengths created an anti-tumor microenvironment, marked by higher populations of naïve macrophages, M1 macrophages, and activated CD8+ T cells (Figure 1B, row (iii), C, row (iii), and F, row (iii)). This was accompanied by higher IFN-γ levels, indicating an enhanced anti-tumor immune response (Figure 1G and H). The secretion of IFN-γ by activated CD8+ T cells played a crucial role in promoting the differentiation of naïve macrophages into the M1 state. Notably, the length of the cell cycle did not significantly affect IL-4 levels.

### The time courses of the population sizes indicate relationships between cell types

In addition to investigating how TME-specific model parameters influence tumor dynamics, we explored the relationships between TME components. We observed a strong similarity between the dynamics of cancer cells and the concentrations of IL-4 in the TME (**Figure 2A, I, and J**).

**Fig 2.**
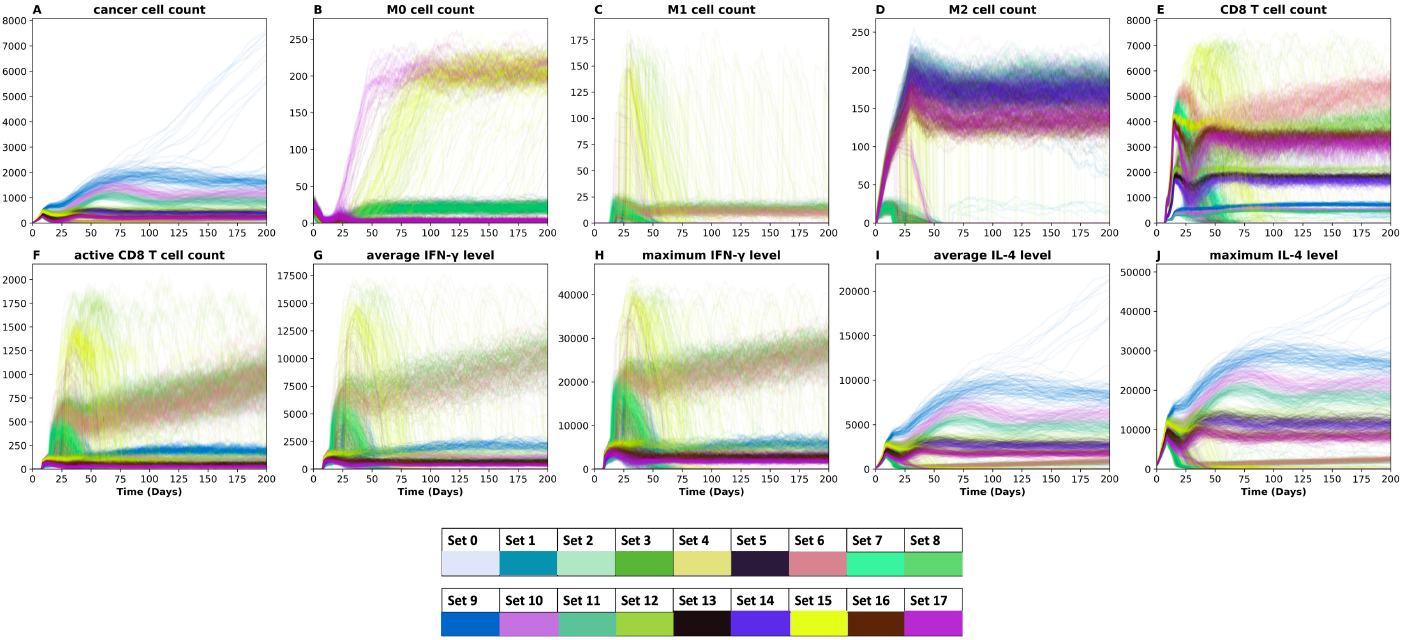
Predicted dynamics demonstrate relationships between different cell types. A) cancer cell count. B) M0 cell count. C) M1 cell count. D) M2 cell count. E) CD8 T cell count. F) active CD8 T cell count G) average IFN-γ level. H) maximum IFN-γ. I) average IL-4 level. J) maximum IL-4 level. All 100 simulations for each parameter set are shown, throughout the 200-day time course.

Both the average and maximum levels of IL-4 exhibited similar patterns to the number of cancer cells, indicating a close relationship between IL-4 and tumor activity. Notably, tumor cells themselves secrete IL-4, directly influencing the concentration of IL-4 within the TME and contributing to the pro-tumor effects. The dynamics of active CD8+ T cells mirrored those of IFN-γ, including both average and maximum IFN-γ levels (**Figure 2F, G, and H**). This correspondence highlights the intricate interactions within the TME, where fully activated T cells secrete IFN-γ, promoting differentiation of naïve macrophages to the M1 state. In contrast, the dynamics of macrophages did not exhibit any distinct trends in relation to the other cell population counts or cytokine levels, suggesting complex and context-dependent behavior.

Similarly, the total CD8+ T cell count did not display a clear trend, indicating their population size is also independent of other TME factors. These findings provide valuable insights into the dynamic interplay between different cell populations within the TME, highlighting the significant roles played by IL-4 and IFN-γ in influencing tumor behavior and the activity of immune cells.

The dynamics of pairs of TME components reveal relationship between cancer cells and active CD8+ cells, underscoring their interconnected dynamics within the TME. **Figure 3** shows the dynamic relationships between cancer cells and every other cell type and diffusible factor in the model. We plot the time courses for the 100 replicates of the model, averaged at each time point, for the immune cell recruitment rates and cancer cell cycle lengths considered above (**Figure 1**). Thus, there are 18 curves represented in **Figure 3**. In the final state (darkest points), a noticeable divergence is observed in the relationship between cancer cells and other cell types, particularly active CD8+ T cells, suggesting distinct tumor states. Most prominently, two states emerge from our simulations (**Figure 3E**): a pro-tumor state characterized by a low count of active CD8+ T cells and a high count of cancer cells, and an anti-tumor state marked by a high count of active CD8+ cells and a low count of cancer cells. This dichotomy highlights the opposing effects of CD8+ T cells on tumor growth, indicating the pivotal role of T cells in shaping the tumor microenvironment. Since the activation of CD8+ T cells is dependent on IFN-γ, there is also a relationship between the cancer cell count and IFN-γ levels (**Figure 3F and G**). We again see two clear regimes: low IFN-γ and high cancer cell count; and high IFN-γ with few cancer cells. Thus, there is a tight link between cancer cells, CD8+ T cells, and IFN-γ.

**Fig 3.**
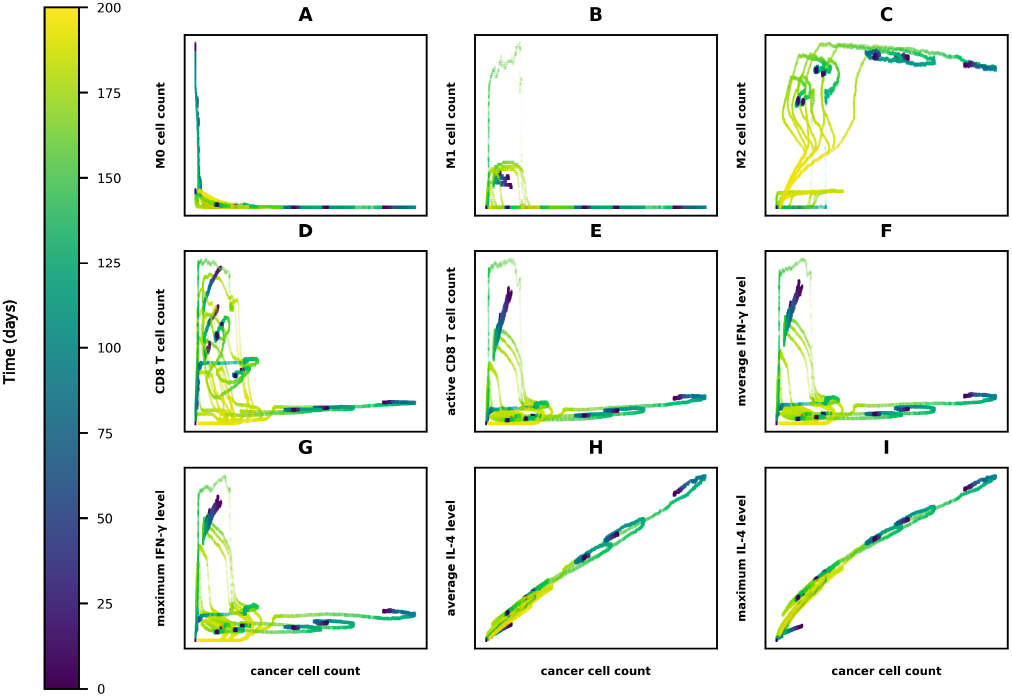
Dynamic relationships present between cancer cells and components of the TME. Cancer cell count versus A) M0 cell count, B) M1 cell count, C) M2 cell count, D) CD8 T cell count, E) active CD8 T cell count, I) average IFN-γ level, J) maximum IFN-γ level, H) IL-4 level, and I) maximum IL-4 level.

Furthermore, we find a significant relationship between M2 macrophages and the population of cancer cells (**Figure 3C**). Initially, as the M2 macrophage population increases, the size of the cancer cell population also increases. This makes sense, as cancer cells influence macrophages through the secretion of IL-4, which promotes the transition of naïve macrophages to M2 macrophages. M2 macrophages also secrete IL-4, creating a positive feedback loop that further reinforces their influence on the cancer cell population. Thus, we also see a direct relationship between the cancer cell trajectories and IL-4 levels (**Figure 3H,I**). The number of cancer cells increases proportionally as the IL-4 concentration increases.

Interestingly, we observe a point in the simulation where the cancer cell population begins to decline, accompanied by a decrease in the M2 macrophage population. This downward trajectory suggests that other factors beyond the relationship between M2 macrophages and cancer cells come into play, potentially influencing the dynamics of the tumor microenvironment. These findings highlight that the mutual influence between M2 macrophages and cancer cells, mediated by IL-4, contributes to the dynamics of the tumor microenvironment.

### The tumor end state can be characterized by the ratio of cancer cells to T cells

The interaction between cancer cells and active CD8+ T cells emerges as a critical determinant of the final state within the tumor microenvironment. We investigated the relationship between the numbers of cancer cells and CD8+ T cells, where two distinct regimes can be discerned (**Figure 4A**). We defined a favorable anti-tumor state to be where the number of CD8+ T cells exceeds the number of cancer cells (yellow region in **Figure 4A**). Similarly, an unfavorable pro-tumor state occurs when the number of cancer cells is greater than the number of CD8+ T cells (purple region in **Figure 4A**). Thus, the threshold between these states is an equal number of cancer cells and CD8+ T cells (**Figure 4A**, dashed black line). We further focused on the final tumor state (**Figure 4B**) to study how particular cell populations and diffusible factors influence the tumor end-state. Here, we display the final number of active CD8+ T cell and cancer cell for all 100 replicates of the 18 parameter combinations shown in **Figure 4A**. The yellow data points represent the anti-tumor state, indicating a robust immune response, with active CD8+ T cells effectively suppressing tumor growth. Conversely, the purple data points represent the pro-tumor state, suggesting a suppressed immune state and creating a favorable environment for tumor growth to prevail. The contrasting pro- and anti-tumor states highlight the significance of the interplay between cancer cells and active CD8+ T cells in shaping the overall dynamics and outcomes of the TME.

**Fig 4.**
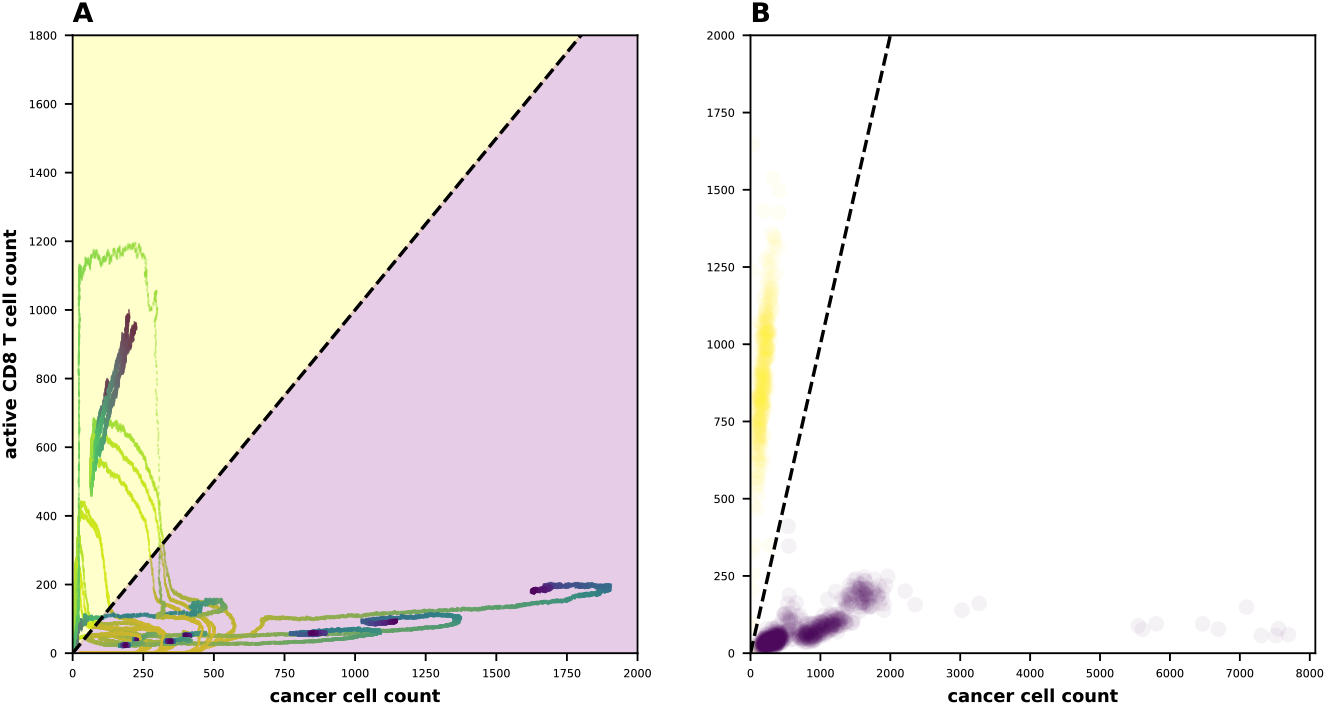
Ratio of cancer cell to active CD8 cells defines final state of the TME. The color scheme differentiates two distinct end-states based on the relationship between these cell populations. The yellow data points represent the anti-tumor state, characterized by a low cancer cell count and a high count of active CD8 cells. The purple data points represent the pro-tumor state, exhibiting a high cancer cell count and a low count of active CD8 cells.

### Regression analysis quantifies components of the TME influence the tumor end state

We performed regression analysis to determine how specific components of the TME (naïve, M1, and M2 macrophages, and average and maximum levels of IL-4 and IFN-γ) influence the predicted final tumor state (ratio of cancer cells to active CD8+ T cells). We performed cross-validation to quantify the predictive ability of the regression analysis. The training accuracy was greater than 0.85 for all test models (Supplementary Figure S1), indicating this analysis can be used to quantify the relationship between TME components and the predict tumor end state. We use the VIP score for each component to quantify how strongly a TME component influences the end state (**Figure 5A**), where a VIP score greater than one indicates a strong influence on the final tumor state. Furthermore, we consider the direction in which the TME component shifts the end state. Here, the loadings of each cell population were examined to assess their contributions to the tumor microenvironment at specific time points and their association with the tumor microenvironment state. The resulting **Figure 5B** displays these loadings, with yellow bars representing variables associated with an anti-tumor state and purple bars indicating variables associated with a pro-tumor state. Notably, variables with a VIP score above 1, indicated by bolded bars, were considered to have significant importance in the model.

**Fig 5.**
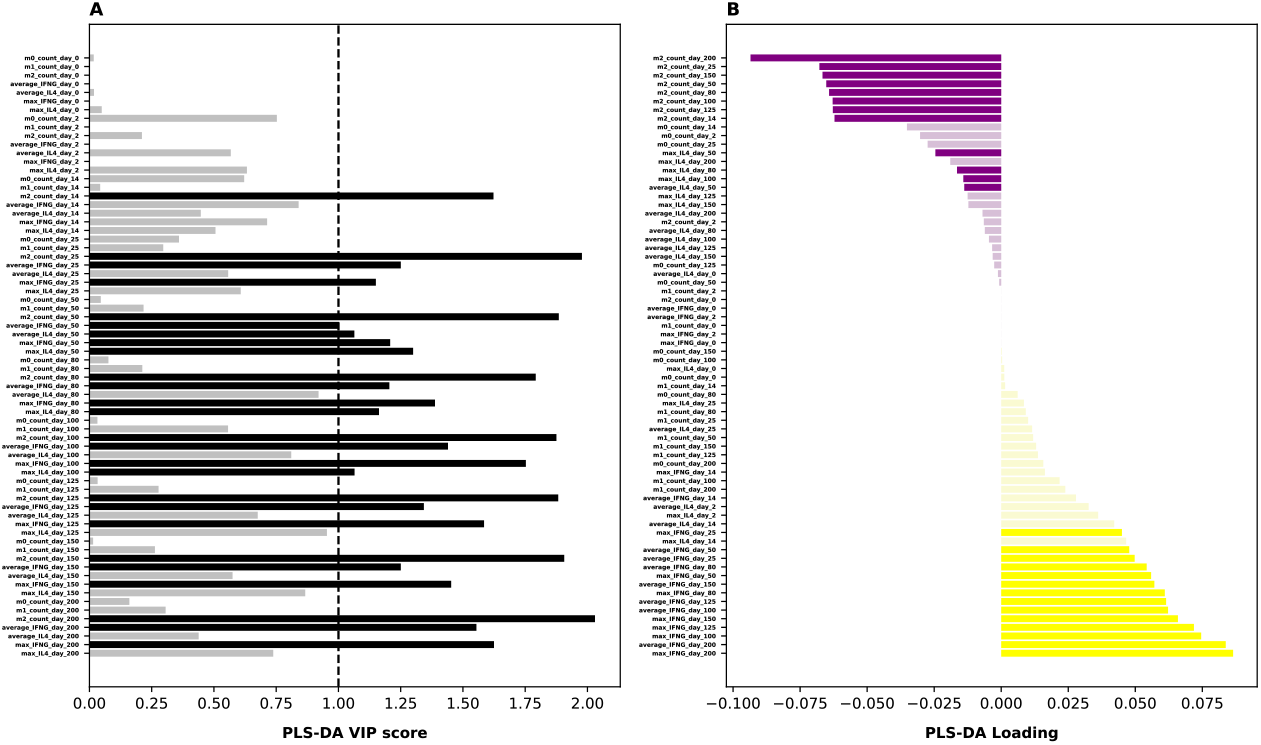
Regression analysis reveals influential factors associated with the final tumor state. A) VIP scores for model inputs from PLS-DA, with black indicating variables that have a VIP score above 1. B) PLS-DA loading values among the cell populations indicates significant influence on the tumor microenvironment. Pro-tumor state indicated by the purple bars and anti-tumor state indicated by the yellow bars. Darker bars indicate inputs with VIP score greater than 1.

Our analysis predicts that at particular time points, the M2 count, average IFN-γ, maximum IFN-γ, and both maximum and average levels of IL-4 influence the end state. We considered the impact of the TME components at 10 distinct time points: day 0, 2, 14, 25, 50, 80, 100, 125, 150, and 200. Prior to day 14, none of the TME components are shown to strongly contribute to the tumor end state. However, starting at day 14 and continuing throughout the 200-day simulation, the number of M2 macrophages is predicted to be a major contributor to the final tumor state (**Figure 5A**). As expected, the negative loading of the M2 count (**Figure 5B**) shows that higher numbers of M2 macrophages shift the tumor towards the unfavorable pro-tumor state. At day 25, the average and maximum IFN-γ levels emerge as significant contributors, promoting an anti-tumor state throughout the remaining time points. Moreover, after day 25, IL-4 appears as a significant contributor, particularly during specific time points. The average and maximum IL-4 levels at day 50 and the maximum IL-4 levels at day 80 are identified as key factors in influencing the number of cancer cells. However, it is important to acknowledge that the influence of IL-4 is not consistent throughout the entire time course. This suggests that IL-4 plays a significant role in certain stages of tumor progression, while its impact may be less pronounced during other times. We further confirm the importance of M2, IL-4, and IFN-γ by plotting their values for each tumor end state (Supplementary Figures S2-S5). Altogether, these findings highlight the influential role of M2 count and IL-4 levels at specific time points in promoting an anti-tumor state, while average and maximum IFN-γ levels promote a pro-tumor state within the TME.

Given the significant role of macrophages in shaping the final tumor state, we investigated the impact of macrophage-based immunotherapies on altering the tumor end state. We simulated three treatments: inhibiting macrophage recruitment to the tumor, depleting macrophages in the tumor, and reeducating M2 macrophages to go to the M1 state. For each case, we ran 100 replicates and analyzed the tumor end state (**Figure 6**). We determined the fraction of end states that are unfavorable (“pro-tumor”: the number of cancer cells exceeds the number of CD8+ T cells) and the fraction of favorable end states (“anti-tumor”: there are more CD8+ T cells than cancer cells). In the absence of any treatment, most of the end states were classified as unfavorable (**Figure 7**), aligning with expectations. However, when evaluating the three different treatments, we found that the reeducation approach, aimed at converting macrophages into an immune-promoting phenotype, emerged as the most effective strategy in promoting an anti-tumor end state. This finding suggests that targeting and reprogramming macrophages holds promise as a therapeutic strategy to shift the tumor microenvironment towards an immune-responsive and anti-tumor state.

**Fig 6.**
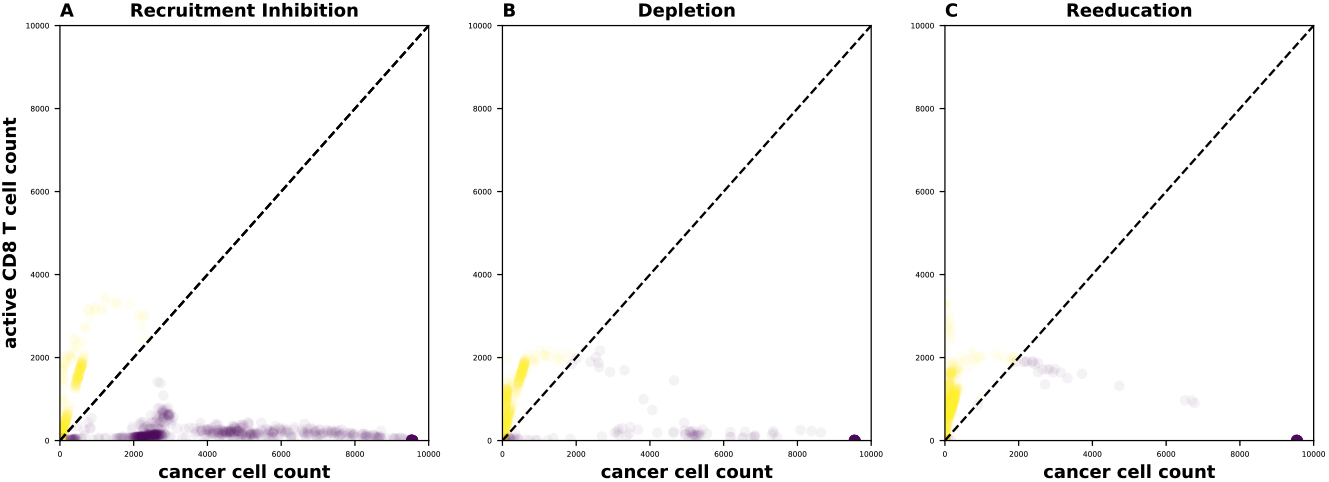
Impact of macrophage-based immunotherapies on tumor end state. A) Recruitment inhibition B) Depletion C) Reeducation. 100 replicates were run for each case and tumor end state was analyzed. Yellow indicates an anti-tumor end state. Purple indicates a pro-tumor end state.

**Fig 7.**
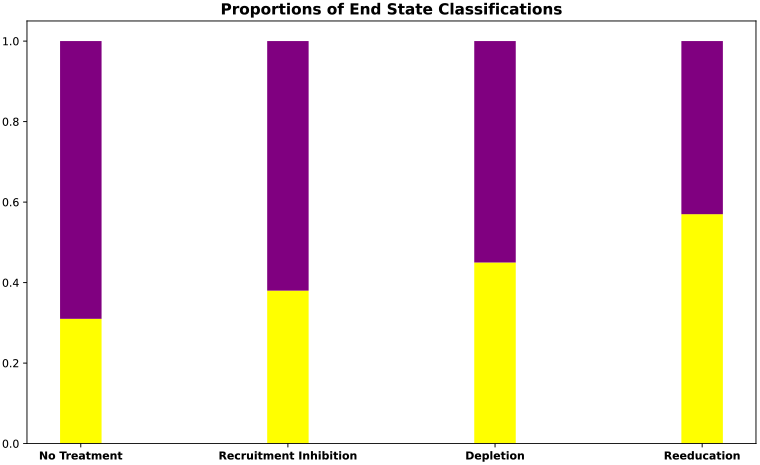
Proportion of pro- and anti-tumor end states after macrophage-based immunotherapy treatments. Yellow indicates an anti-tumor state. Purple indicates the pro-tumor state. Macrophage-based treatment increases the predicted proportion of anti-tumor end state classifications.

## Discussion

In this study, we utilized an ABM to simulate the interactions between tumor and immune cells within the TME. The model incorporated three primary cell types: cancer cells, macrophages (naïve, M1, and M2), and T cells (active CD8+ and inactive). We also considered the production and diffusion of two key diffusible factors, IL-4 and IFN-γ, which play crucial roles in influencing the immune state of the TME. By varying parameters such as cancer cell cycle length and the rates at which macrophages and T cells are recruited, we simulated the dynamic behavior of the tumor and investigated the effects of these cell-specific properties factors on tumor progression. Our analysis revealed the relationships between different cell populations and highlighted the importance of macrophages and T cells in shaping the TME.

It is important to address the concept of pro-tumor and anti-tumor end states within the TME and the factors that contribute to them. Our findings consider that a high number of active CD8+ T cells and low numbers of cancer cells are associated with an anti-tumor end state^19^.

Conversely, a pro-tumor end state is characterized by a low presence of active CD8+ T cells and a high density of cancer cells, indicating a compromised immune response and a supportive environment for tumor growth^19^. Given these definitions, we explored how TME components contribute to these pro-tumor and anti-tumor end states. The presence of macrophages in the TME plays a crucial role, as macrophages can exhibit different phenotypes with opposing effects on tumor progression^20^. M1 macrophages have immune-stimulatory properties and promote an anti-tumor response, while M2 macrophages have immune-suppressive properties and contribute to a pro-tumor environment^21^. Therefore, the balance between M1 and M2 macrophages is critical in shaping the TME and determining the end state. The production and diffusion of key diffusible factors, such as IL-4 and IFN-γ, influence the immune state of the TME as well. IL-4 is typically associated with an immune-suppressive environment, promoting a pro-tumor state, while IFN-γ is linked to an immune-stimulatory environment, favoring an anti-tumor response^22^. The levels and distribution of these factors within the TME can significantly impact the immune cell populations and their interactions, thereby influencing the end state. Our work quantifies the importance of these TME components, at specific times during tumor progression. By establishing the influence of the TME factors, we can use the model to inform strategies to target the influential TME components.

The study demonstrated that macrophage-based immunotherapies, specifically the reeducation approach targeting macrophage phenotype conversion, showed promise in promoting an anti-tumor end state within the TME. By reprogramming macrophages to an immune-promoting phenotype, the study suggests that it may be possible to shift the tumor microenvironment towards an immune-responsive, anti-tumor state. These findings provide valuable insights into the potential of targeting macrophages as a therapeutic strategy for cancer treatment^23^.

This study addresses the complexity of tumor-immune cell interactions and the crucial role played by macrophages in tumor progression. By employing an ABM, the study provides a detailed understanding of the dynamics within the TME and sheds light on the underlying mechanisms that govern tumor behavior and response to immunotherapy. The use of computational models, based on biological experiments, allows for reliable and physiologically-relevant predictions of tumor behavior, which may not be feasible with purely experimental approaches^24^. The study emphasizes the significance of using computational modeling to gain a mechanistic and quantitative understanding of tumor-immune interactions. Furthermore, the study highlights the potential of macrophage-based immunotherapies as a promising avenue for cancer treatment^25^. Overall, our findings contribute to ongoing efforts to develop more effective immunotherapies and personalized treatment strategies for cancer patients^26^.

While the ABM used in this study provides valuable insights into the dynamics of the TME, it is important to acknowledge its limitations. The model represents an idealized 2D representation of the tumor slice, and the simulations were performed without considering the influence of other factors such as angiogenesis, genetic heterogeneity, or the presence of other immune cell types. Therefore, the model may not fully capture the complexity and heterogeneity of the actual tumor microenvironment. Future studies should aim to incorporate these additional factors to obtain a more comprehensive understanding of tumor-immune interactions. Moreover, the parameter values used in the model were derived from previous modeling efforts or based on experimental observations. While this approach provides a basis for reliable predictions, it is essential to acknowledge that the model’s accuracy depends on the accuracy of these parameter values. Future studies may refine these parameters and validating the model predictions using experimental data.

Building upon the insights gained from this study, future research directions can focus on several aspects. Firstly, incorporating additional cell types and factors into the ABM, such as other immune cell populations, cytokines, and the tumor microenvironment’s extracellular matrix, will provide a more comprehensive representation of the TME and enable a deeper understanding of the intricate interactions within it. Secondly, further exploration of macrophage-based immunotherapies is warranted. While the reeducation approach showed promise in promoting an anti-tumor end state, more in-depth investigations are needed to optimize the therapy and assess its efficacy in preclinical and clinical settings. Future studies should consider the development of targeted therapies that selectively modulate macrophage phenotypes to maximize the immune response against tumors while minimizing potential side effects. Finally, combining computational modeling combined with experimental approaches, such as *in vitro* and *in vivo* preclinical studies, can provide a more holistic perspective on tumor-immune interactions. Experimental validation of the model predictions, as well as the integration of patient-specific data, can enhance the accuracy and translational potential of computational models.

## Conclusions

This study highlights the importance of computational modeling in understanding the complex dynamics of tumor-immune interactions within the tumor microenvironment. The findings underscore the potential of macrophage-targeted immunotherapies and provide insights into the underlying mechanisms that govern tumor behavior. While acknowledging the study’s limitations, future research can build upon these findings to refine and expand the models, explore new therapeutic strategies, and ultimately contribute to the development of more effective and personalized cancer treatments.

## Author Contributions

S.D.F. conceived of the study. M.A. carried out computational model simulations. M.A. and N.T. performed analyses. S.D.F. supervised the project and provided financial support. All authors discussed the results and contributed to the final manuscript.

## Declaration of Interests

The authors declare no competing interests.

## Acknowledgements

The authors thank Dr. Colin Cess for guidance in performing model simulations. This work was supported by the USC Summer Oncology Research Fellowship (M.A.), the USC Student Opportunities for Academic Research (N.T.), and the USC Center for Computational Modeling of Cancer.

## Data Availability

No new data was generated as part of this study. The agent-based model used in this work has been previously published and is available publicly.

## Notes

### Competing Interest Statement

The authors have declared no competing interest.

