## Supplementary Material for "Agent-based modeling of tumor-immune interactions reveals determinants of final tumor states"

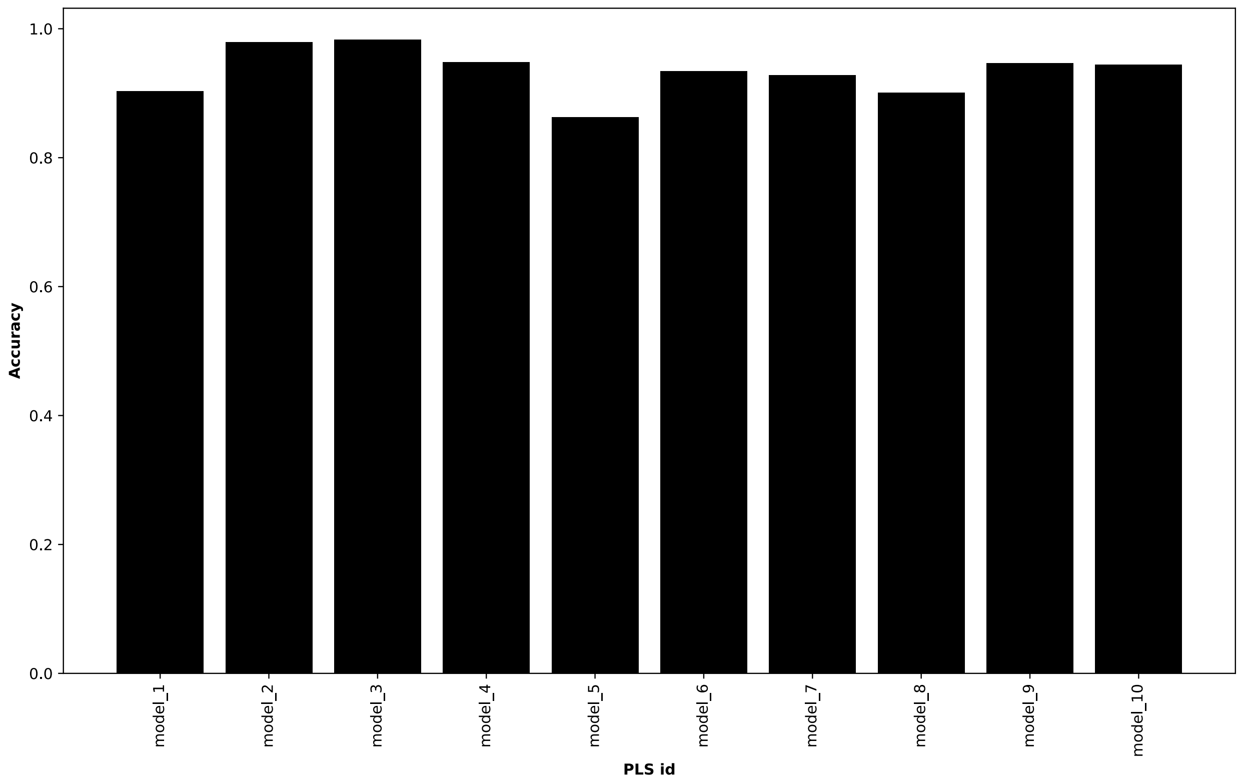


**S1. High accuracy is maintained across all 10 PLSR models.** A suite of 10 PLSR models was generated and cross validated to predict how each model component contributes to the predicted final ratio of tumor cells to active CD8 T cells. All models displayed relatively high accuracy.

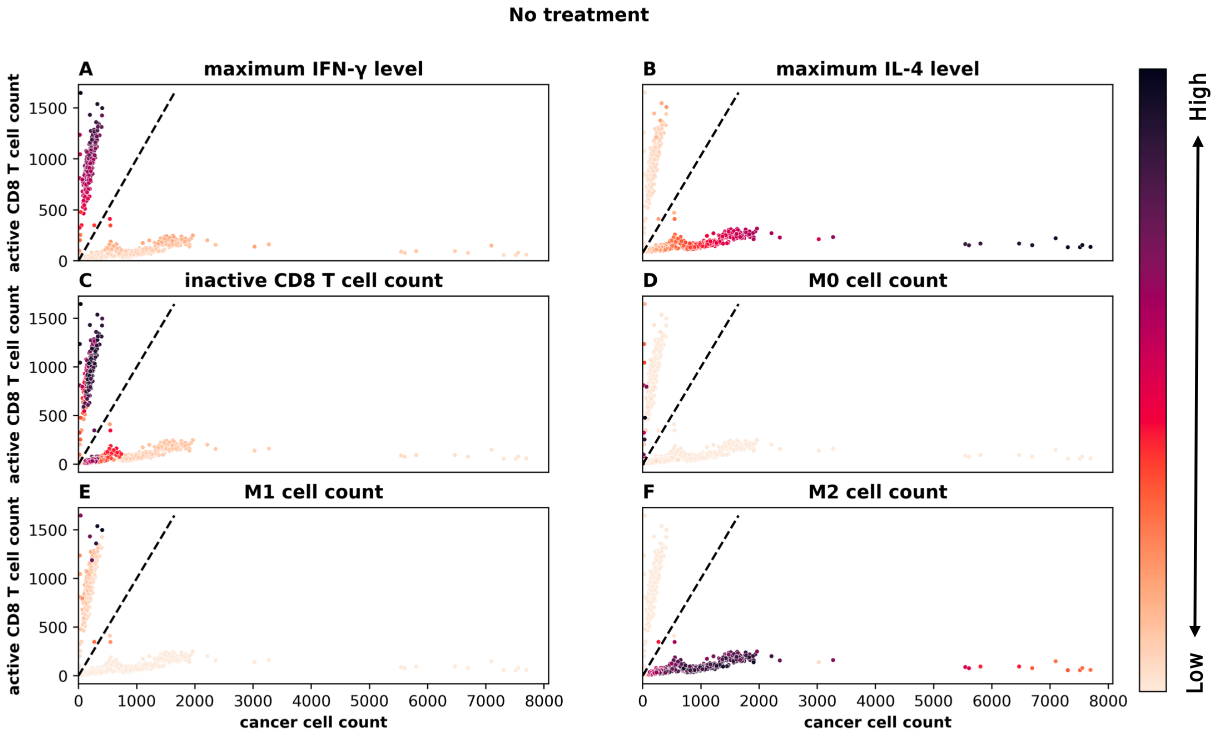


**S2. Predicted final tumor state for no treatment.** A) cancer cell count vs. maximum IFN-γ level. B) cancer cell count vs. maximum IL-4 level. C) cancer cell count vs. CD8 T cell count. D) cancer cell count vs. M0 cell count. E) cancer cell count vs. M1 cell count. F) cancer cell count vs. M2 cell count

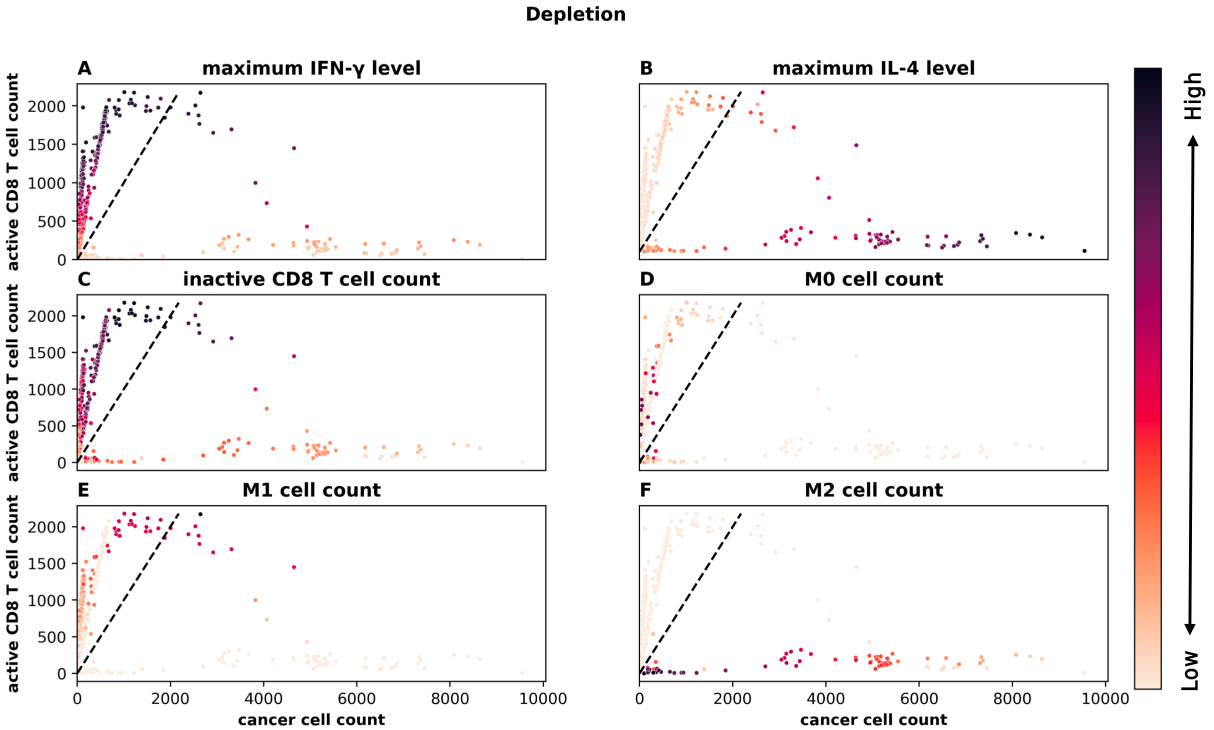


**S3. Impact of macrophage depletion treatment on final tumor state.** A) cancer cell count vs. maximum IFN-γ level. B) cancer cell count vs. maximum IL-4 level. C) cancer cell count vs. CD8 T cell count. D) cancer cell count vs. M0 cell count. E) cancer cell count vs. M1 cell count. F) cancer cell count vs. M2 cell count

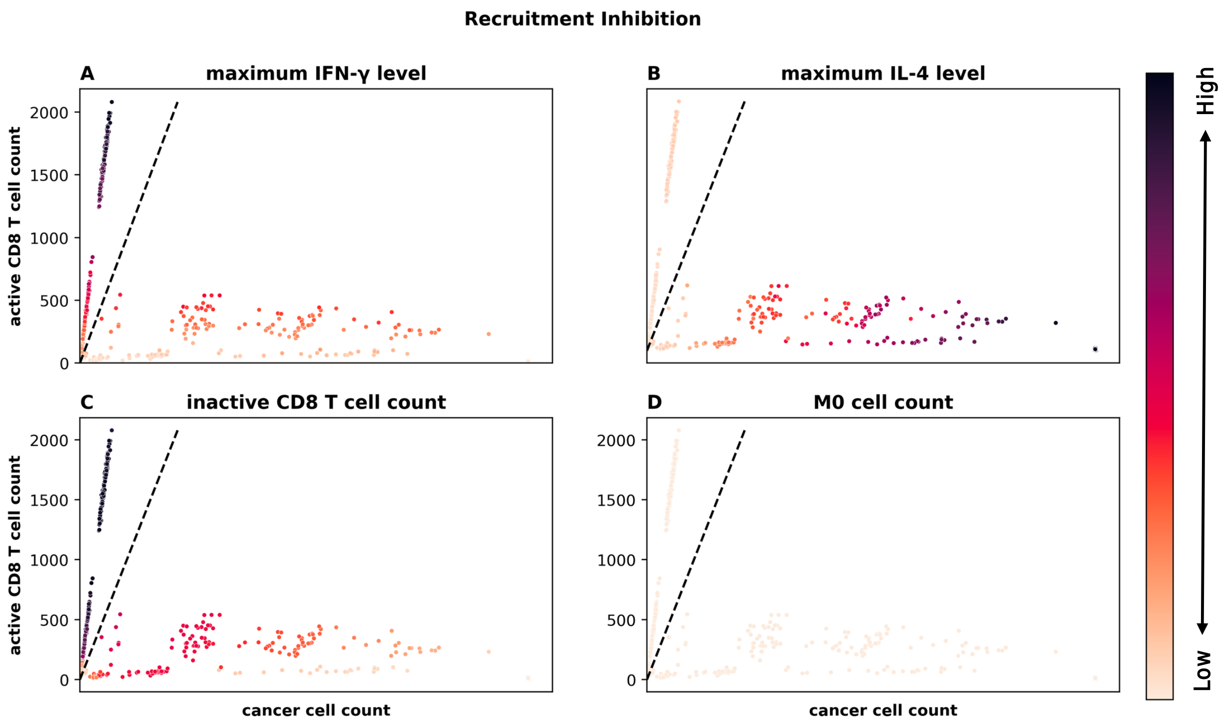


**S4. Impact of inhibition of macrophage recruitment treatment on final tumor state.** A) cancer cell count vs. maximum IFN-γ level. B) cancer cell count vs. maximum IL-4 level. C) cancer cell count vs. CD8 T cell count. D) cancer cell count vs. M0 cell count.

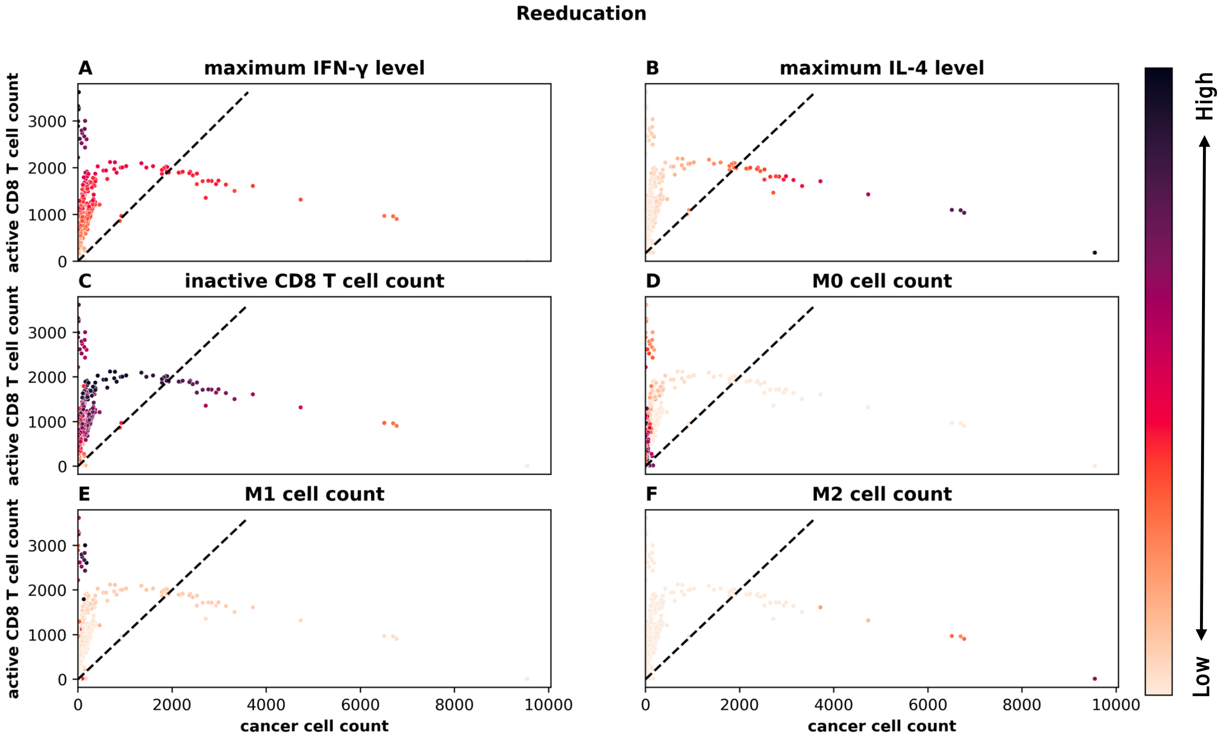


**S5. Impact of macrophage reeducation treatment on final tumor state.** A) cancer cell count vs. maximum IFN-γ level. B) cancer cell count vs. maximum IL-4 level. C) cancer cell count vs. CD8 T cell count. D) cancer cell count vs. M0 cell count. E) cancer cell count vs. M1 cell count. F) cancer cell count vs. M2 cell count
